# Impact of trait exaggeration on sex-biased gene expression and genome architecture in a water strider

**DOI:** 10.1101/2020.01.10.901322

**Authors:** William Toubiana, David Armisén, Corentin Dechaud, Roberto Arbore, Abderrahman Khila

## Abstract

Exaggerated secondary sexual traits are widespread in nature and often evolve under strong directional sexual selection. Although heavily studied from both theoretical and empirical viewpoints, we have little understanding of how sexual selection influences sex-biased gene regulation during the development of sex-specific phenotypes, and how these changes are reflected in genomic architecture. This is primarily due to the lack of a representative genome and transcriptomes to study the development of secondary sexual traits. Here we present the genome and developmental transcriptomes, focused on the legs of the water strider *Microvelia longipes*, a species where males exhibit strikingly long third legs used as weapons. The quality of the genome assembly is such that over 90% of the sequence is captured in 13 scaffolds. The most exaggerated legs in males were particularly enriched in sex-biased genes, indicating a specific signature of gene expression in association with sex-specific trait exaggeration. We also found that male-biased genes showed patterns of fast evolution compared to non-biased and female-biased genes, indicative of directional or relaxed purifying selection. Interestingly, we found that female-biased genes that are expressed in the third legs only, but not male-biased genes, were over-represented in the X chromosome compared to the autosomes. An enrichment analysis for sex-biased genes along the chromosomes revealed that they can arrange in large genomic regions or in small clusters of two to four consecutive genes. The number and expression of these enriched regions were often associated with the exaggerated legs of males, suggesting a pattern of common regulation through genomic proximity in association with trait exaggeration. Our findings shed light on how directional sexual selection drives sex-biased gene expression and genome architecture along the path to trait exaggeration and sexual dimorphism.

## Introduction

Sexual dimorphisms, or phenotypic differences between males and females of the same species, is one of the most common sources of phenotypic variation in nature ^1,2^. Understanding how this process is regulated in a sex-specific manner at the genomic level, despite the shared genome between males and females, still poses an important challenge ^3^. Differences in gene expression have emerged as a common mechanism to explain phenotypic differences among individuals sharing almost the same genome ^4,5^. In the last decade, a large number of studies have characterized genes with sex-biased expression in a variety of species, leading to an emerging framework attempting to link sex-biased gene expression to phenotypic divergence of the sexes ^4–6^. However, these studies have mostly focused on adult gonads or whole-body transcriptomic datasets, which are unsuited to understand how secondary sexual characters are built during development ^4,5^. Among the countless examples of sexual dimorphism, some species have evolved extreme patterns whereby males, generally, develop such drastic phenotypes that they appear exaggerated compared to homologous traits in the other sex or to other body parts ^7–10^. These growth-related sexual traits have received lots of attention in developmental genetics, but we still lack a general understanding of the genomic regulation underlying their development ^7,11–21^.

We aimed here to assess the effect of sexual selection on the sex-specific regulation of gene expression and genome architecture in the water strider *Microvelia longipes* (Heteroptera, Gerromorpha, Veliidae), an emerging model in the field of sexual selection and trait exaggeration ^22^ (see also companion paper Toubiana et al.). *M. longipes* is a hemimetabolous insect that displays a striking case of sex-specific exaggerated trait where some males develop extremely long rear legs compared to females. The length of the rear legs (third legs) in males is under strong directional sexual selection and these legs are used as weapons to kick opponents away from the sites where females mate and lay eggs ^22^. Such directional selection is associated with the evolution of disproportionate growth (i.e. hyperallometry) in male rear legs. Here we study the genomic regulation underlying the elaboration of this exaggerated phenotype in order to shed light on the role of sexual selection in shaping genome evolution. We generated a high-quality genome of *M. longipes*, with chromosome-scale resolution, and compared the expression, molecular evolution and genomic location of sex-biased genes in the three pairs of legs at a developmental stage where the legs diverge between the sexes (companion paper Toubiana et al.). Combined, our approach first identified signatures of trait exaggeration in terms of sex-biased gene expression patterns and sequence evolution. Second, it characterized chromosomes and genomic regions that are enriched in the sex-biased genes associated with the directional sexual selection applying to male exaggerated legs in *M. longipes*.

## Results

### *De novo* assembly and automatic annotation of *M. longipes* genome

To study the genetic mechanisms underlying extreme growth of male legs, we generated *de novo* the genome of *M. longipes* (Figure 1A) using lines established from a French Guiana population that were inbred through 15 sib-sib crosses ^22^. Next generation sequencing and k-mer frequency distribution in raw sequencing reads estimated *M. longipes* genome size to about 0.67Gb (Supplementary table 1; see material and methods). Genome assembly combined multiple mate-pair Illumina libraries, PacBio, and Dovetail Hi-C/Hi-Rise libraries ^23–25^ (Supplementary table 1; see Material and Methods). The final assembly generated chromosome-length scaffolds with scaffold N50=54.155 Mb and contig N50=216.72 Kb (Supplementary table 1). Over 90% of *M. longipes* genome is represented in the thirteen largest scaffolds (Supplementary table 1; see material and methods).

**Figure 1:**
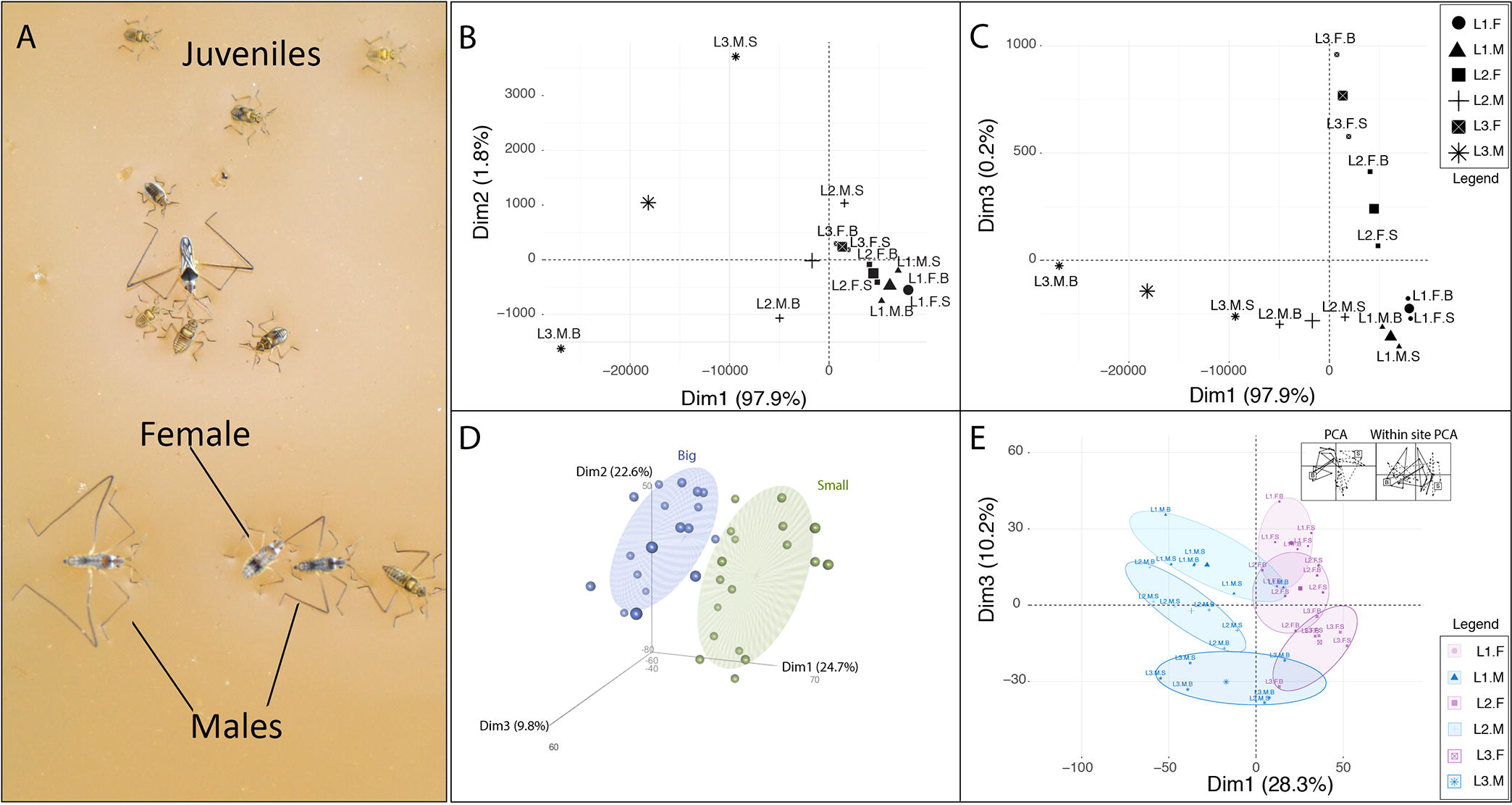
Association between variation in leg length and variation in gene expression. (**A**) *Microvelia longipes* in the wild. (**B-C**) Principal Component Analysis (PCA) on male and female leg length from the long-leg and short-leg selected inbred populations ^22^. (**B**) The first PCA (Dim1) explains primarily differences between legs of the same sex while the second PCA (Dim 2) explains differences between inbred populations, specifically in males. (**C**) The third PCA (Dim 3) explains the differences between sexes. (**D-E**) Principal Component Analysis (PCA) on the whole transcriptomic dataset. (**D**) The three first PCAs (Dim1, 2, 3) recapitulate the variance between the Big (blue) and Small (green) lines. (**E**) Within-Class analysis after correcting for line effects. Dimension 1 separates sexes while Dimension 3 separates legs. The inset represents the Within-Class correction for the line effects.

We then used automatic genome annotation, supported by *de novo* transcriptome-based gene models, to build the gene set of *M. longipes* (see material and methods). This analysis predicted 26 130 genes and 27 553 transcripts. BUSCO analysis, based on the 2018 insect dataset ^26^, revealed that 96% of gene models are present; among these 92% are complete, 3% are fragmented and 1% are duplicated (Supplementary figure 1). We therefore conclude that *M. longipes* genome is more complete than most available insect genomes ^27^.

### Variation in gene expression explains differences in leg length

In *M. longipes*, the most obvious difference between legs is reflected in their size (Figure 1B) ^22^. Principal component analysis (PCA) of adult male and female leg length from two isogenic lines, selected for differences in absolute leg length ^22^, revealed that the first major component of variation encompassed 97% of the total variation and separated samples based on serial homolog in both sexes, with male third legs covering most of the divergence (Figure 1B). The second and third major axes of variation discriminated males from the two lines and sexes, respectively, although they only contributed 2% to the total variation (Figure 1B-C).

To test whether these phenotypic differences correlate with variation in gene expression, we sequenced the leg transcriptomes of males and females from these lines at the 5^th^ nymphal instar – the developmental stage where we observed a burst of growth (See companion paper, Toubiana et al.). If a strict correlation existed between leg length and gene expression, we should predict samples to cluster by serial homolog, then by line, and finally by sex. Instead, the three first major axes of variation in the leg transcriptomes clustered samples based on line (Figure 1D). The line effect accounted for about 60% of the total variation in gene expression, thus potentially hiding signals associated with differences between legs and sexes. We therefore corrected for this line effect, using within class analysis, and generated a new PCA that now separates the sexes in PC1 (28.3% of variation in gene expression), and the serial homologs of the same sex in PC3 (10.3% of the variation in gene expression) (Figure 1E). We conclude that the three main components involved in leg length variation were retrieved in our transcriptome datasets. Yet, conversely to leg morphologies, homologous legs from the two sexes are now more divergent in terms of gene expression than serial homologs from the same sex. We hereafter focus on the effect of sex on gene expression as it represents a major factor underlying leg exaggeration through differences in allometric coefficients ^22^ (Supplementary figure 2).

### Leg exaggeration and sex-biased gene expression

The legs of *M. longipes* males and females differ in their scaling relationships and degree of exaggeration ^22^ (Figure 1A-C, Supplementary figure 2). To determine more specifically the patterns of gene expression underlying the observed sexual dimorphism in scaling relationships, we compared expression profiles of homologous legs between the sexes. All three legs consistently showed over twice as many female-biased than male-biased genes, with the third legs having the highest total number of sex-biased genes (Figure 2A-C). Interestingly, the average degree of differential expression (log2 Fold Change) in male-biased genes correlated with the patterns of leg growth such that the second and the third legs, which are hyper-allometric, showed higher degree of male-biased expression than the first leg, which is iso-allometric (Figure 2D). This pattern was consistent with the general overexpression of male-biased genes in the two exaggerated legs, especially the third, compared to the first legs (supplementary figure 3). This correlation was however absent for female-biased genes (Figure 2D; supplementary figure 3). More than two thirds of the male-biased genes in the first and second legs were shared with the most exaggerated legs, whereas about two thirds of male-biased genes in the third leg were not shared with the other legs (Figure 2E). A hierarchical clustering analysis separated males’ third legs from the other serial homologs in both sexes, confirming that on average the sexual dimorphic expression of these 354 genes is restricted to the most exaggerated leg (Figure 2 F). Furthermore, the third legs showed a high number of female-biased genes, despite the lack of exaggeration in females, suggesting that the development of this extreme sexual dimorphism may also result from the active regulation of specific genes in female’s third legs or their active repression in male’s third legs. Altogether, these results show that the exaggerated third legs of males display unique patterns of sex-biased genes, in terms of number and/or levels of expression, compared to the two other serial homologs.

**Figure 2:**
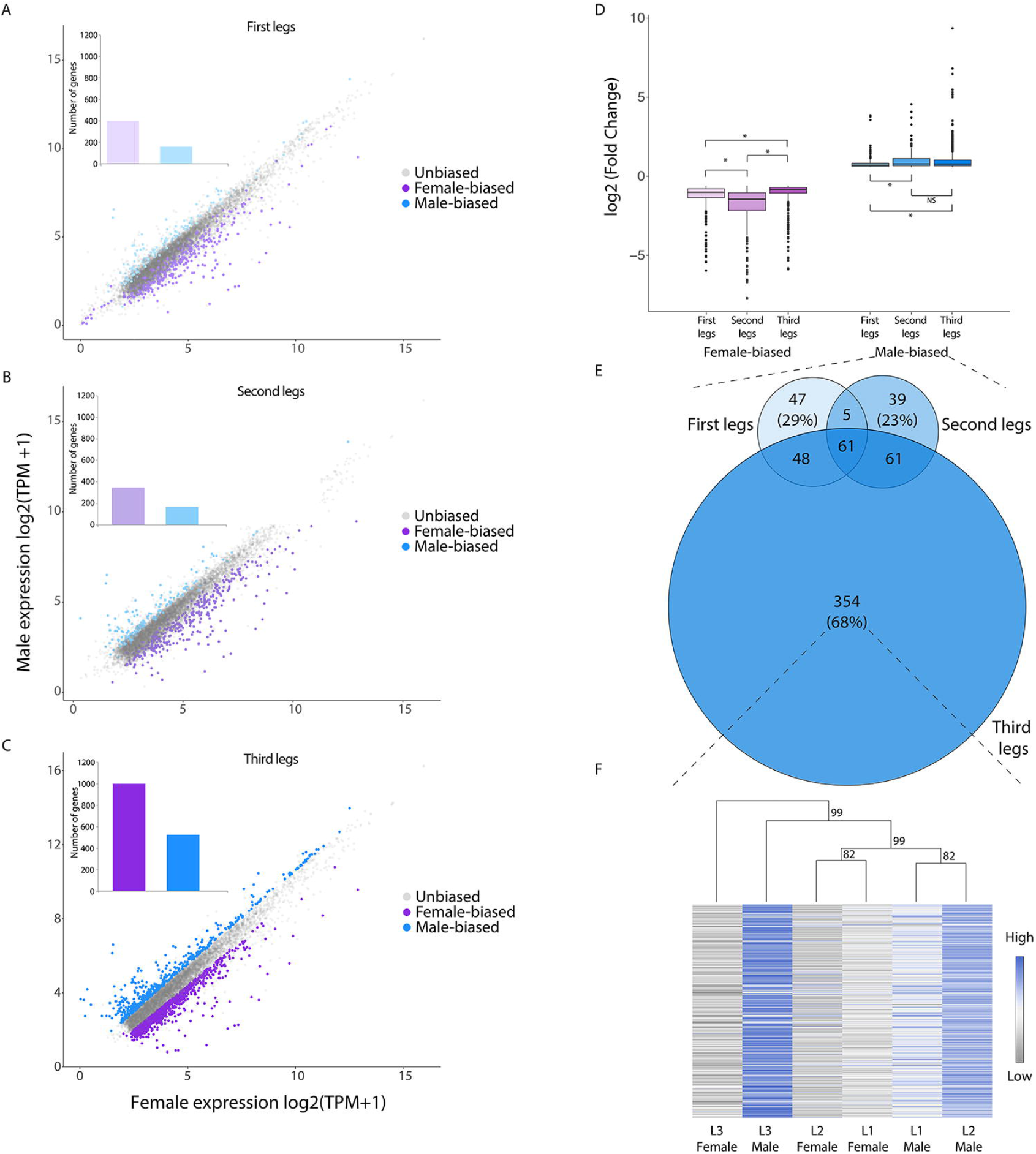
Signature of trait exaggeration among sex-biased genes. (**A-C**) Comparison of gene expression (log2 TPM+1) in male and female legs. Dots highlighted in purple and blue represent genes with significant difference in expression in females and males respectively. Insets indicate the number of female- and male-biased genes in each leg. (**D**) Differences in fold change (Wilcoxon tests) among the sex-biased genes identified in the three pairs of legs independently. (**E**) Venn-diagrams of the male-biased genes identified in the three pairs of legs. Size of the diagrams is proportional to the total number of genes. (**F**) Hierarchical clustering (1000 bootstraps) and heatmap based on average leg expression in males and females for the genes with significant male-biased expression specifically in the third legs of males (n=354).

### Male exaggerated legs are enriched in both leg- and sex-biased genes

Theory predicts that pleiotropy may constrain the evolution of sex-biased gene expression ^28^. Our analysis of sex-biased genes reports a possible crosstalk between tissue and sex regulations in the context of trait exaggeration, which in turn may relax genetic constraint on sex-biased genes. Alternatively, sexual dimorphism may result from post-transcriptional regulation, possibly resulting in rather broad expression of sex-biased genes ^29^. To test these hypotheses, we combined our list of leg-biased genes (genes differentially expressed between the legs of the same sex, see companion paper Toubiana et al.) with the list of sex-biased genes (genes differentially expressed between the same leg in males and females), and performed a four-dimension comparison of fold-change between first and third legs as serial homologs in the Y-axis and males and females in the X-axis (Figure 3).

**Figure 3:**
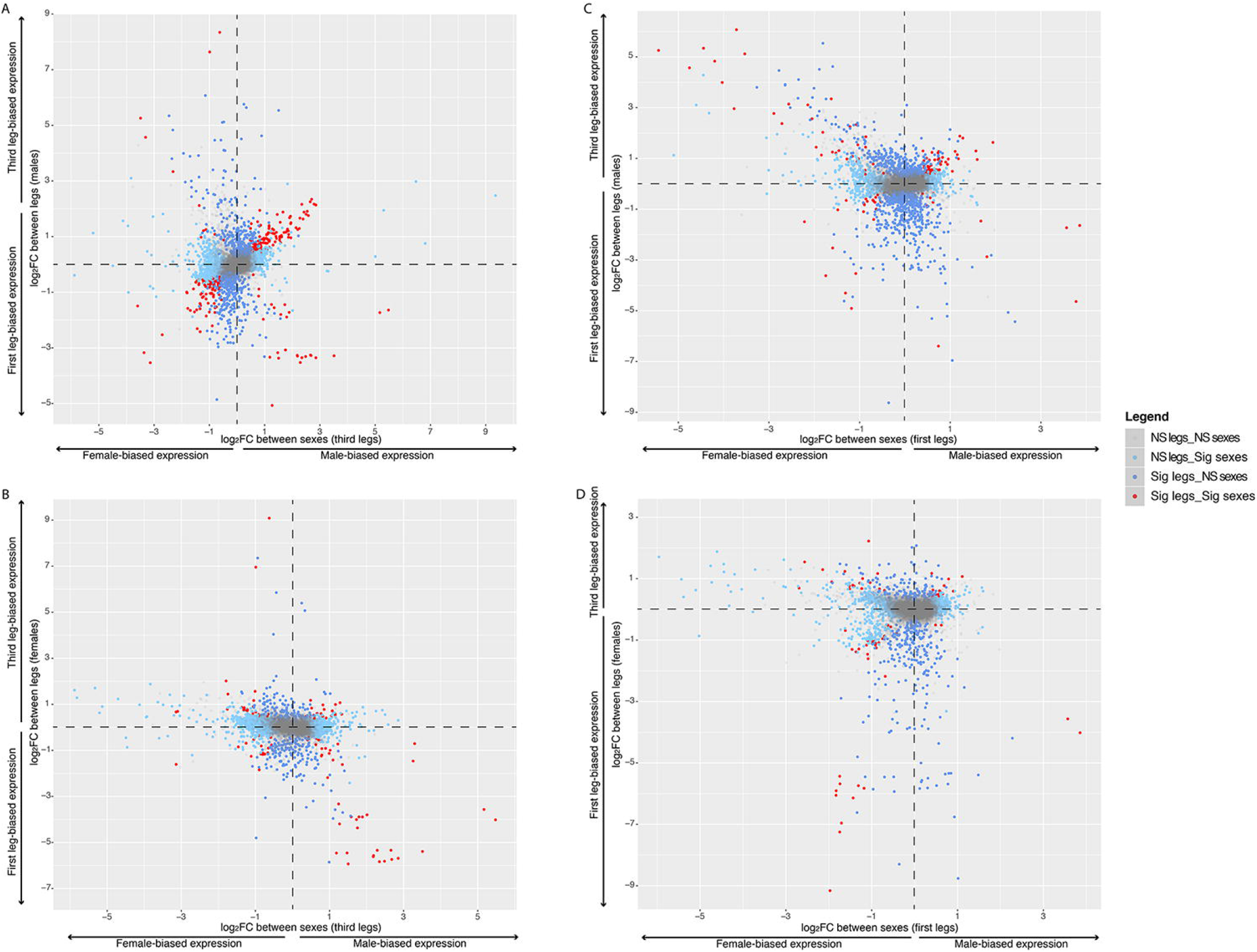
Crosstalk between leg- and sex-biased genes. Light blue dots are sex-biased genes (FC > 0, Padj < 0.05), dark blue dots correspond to leg-biased genes (FC > 0, Padj < 0.05) and red dots are genes with both leg- and sex-biased expressions. For the leg-biased genes, we only illustrated the comparisons between the third and first legs. Data on the second legs are shown in Supplementary figure 4. “NS” indicates, “Non Significant” and “Sig” indicates “Significant”. (**A**) Comparison between sex-biased genes in the third legs and leg-biased genes in males. (**B**) Comparison between sex-biased genes in the third legs and leg-biased genes in females. (**C**) Comparison between sex-biased genes in the first legs and leg-biased genes in males. (**D**) Comparison between sex-biased genes in the first legs and leg-biased genes in females.

Interestingly, genes that are up regulated in male compared to female third legs also tend to be up regulated in male third legs compared to first legs (Fisher’s exact test; p-value < 0.05) (Figure 3A). Conversely, female-biased genes in the third legs tend to be down regulated in males’ third legs compared to the first legs (Fisher’s exact test; p-value < 0.05) (Figure 3A). These patterns were however absent when we analyzed the genes that are differentially regulated in female serial homologs (Fisher’s exact tests; p-values > 0.05) (Figure 3B). Likewise, these associations were lost when we looked at sex-biased genes in the first legs. Male-biased genes in the first legs tend to be under-represented among up regulated genes in male first legs (Fisher’s exact test; p-value < 0.05) (Figure 3C). Moreover, female-biased genes did not show any leg-biased tendency in males (Fisher’s exact test; p-value > 0.05) (Figure 3C). We obtained similar results when we selected leg-biased genes in females, except for the up-regulated genes in the first legs that tend to be female-biased (Fisher’s exact test; p-value < 0.05) (Figure 3D).

In the second legs, which are mildly exaggerated in males, we also recovered enrichment between leg-biased genes in males and sex-biased genes in the second legs, though with fewer genes than the third legs (Fisher’s exact tests; p-value < 0.05) (Supplementary figure 4). Overall, we found an enrichment of genes with both leg- and sex-biased expression that was particularly higher in male exaggerated third legs. This crosstalk highlights possible modularity by which leg-biased genes would have more easily acquired sex-biased expression to promote the exaggerated growth of male third legs without affecting other organs.

### Male-biased genes evolved fast

We have shown that the pattern of sex-biased expression in *M. longipes* legs correlated in several aspects with the elaboration of the exaggerated third legs in males. In several species, sex-biased genes display higher rate of evolution compared to unbiased genes but relatively little is known about their sequence evolution in the context of trait exaggeration. We classified expressed genes in *M. longipes* legs based on their sex-biased expression pattern (male-biased, female-biased and unbiased, respectively) and compared their sequence evolution with four other *Microvelia* species (Figure 4; Supplementary table 2). We generally found that male-biased genes evolved faster than female-biased and unbiased genes; female-biased genes being the slowest evolving (Figure 4; Supplementary table 2). This pattern was true for all species-pair comparisons and remained consistent across legs or when we separated X-linked genes from autosomal genes (Figure 4; Supplementary table 2). Conversely to what has been observed in several organisms, we failed to detect any pattern of faster-X evolution (Supplementary table 2).

**Figure 4:**
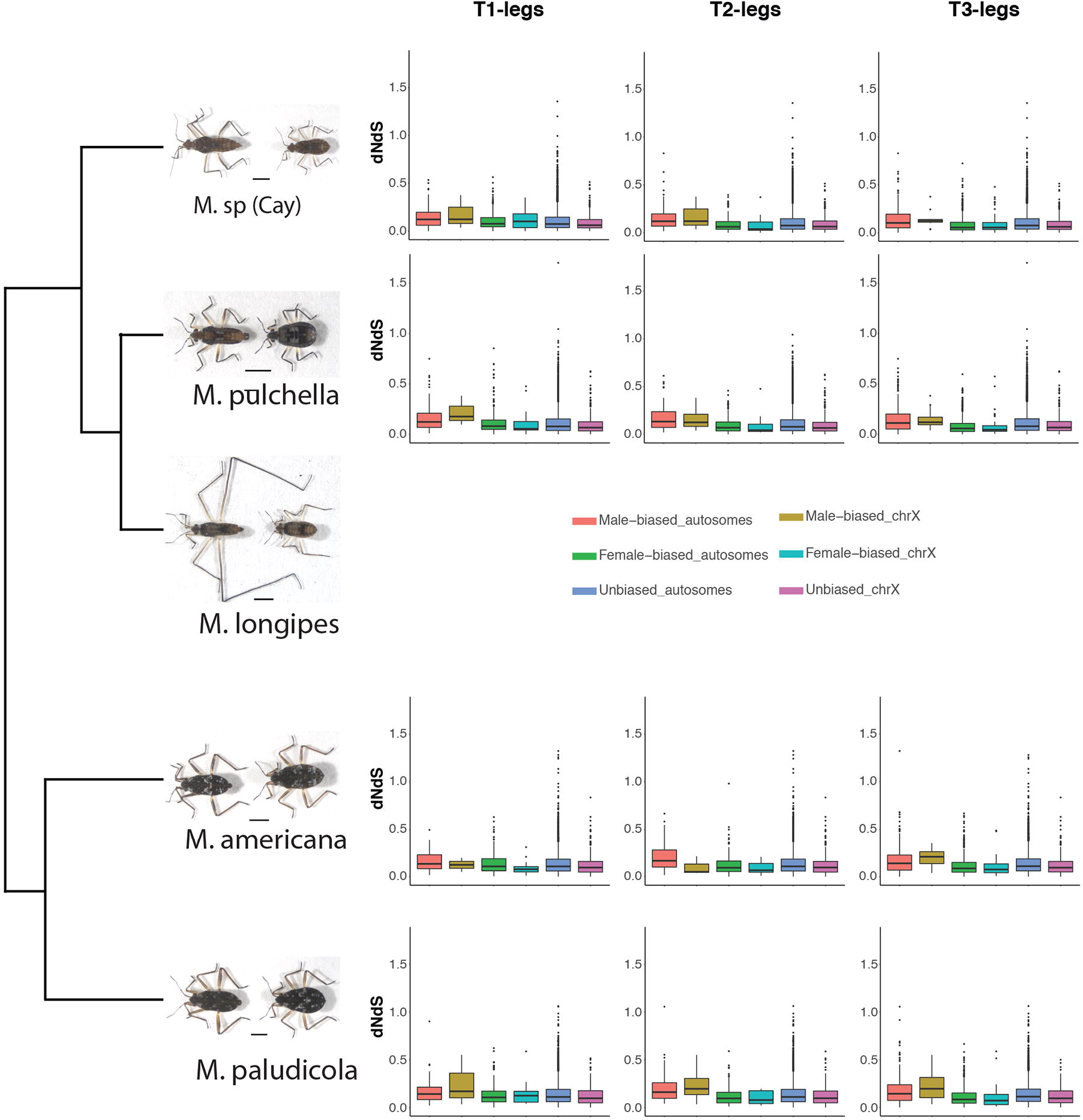
Sequence evolution of genes expressed in *M. longipes* legs. We categorized genes based on their expression profile and genomic location. Boxplots indicate the pair-wise comparisons of dNdS across the three legs between *M. longipes* and each of the four other *Microvelia* species. Statistical analyses are shown in Supplementary table 2.

### Sex-biased gene expression and genome architecture

Theory predicts that sexual selection can be an important driver of genome evolution ^30,31^, and we sought to test this prediction by analyzing the distribution of sex-biased genes along the genome of *M. longipes.* First, we identified the scaffold that corresponds to the X chromosome (see material & methods). Interestingly, our analysis detected enrichment in the X chromosome with female-biased genes of the third legs, but not the two other legs, compared to the autosomes (Figure 5A). The percentage of female-biased genes between the X chromosome and the autosomes in the different legs confirmed that the enrichment observed was caused by an accumulation on the X chromosome of genes specifically biased in the third legs of females (Figure 5A). In contrast, we did not find any significant under- or over-representation of male-biased genes from any of the three legs on the X chromosome (Figure 5A). Because of the known effect of dosage compensation on the expression of genes located on the X chromosome ^4,32–34^, we compared the levels of expression of all X chromosome genes between the sexes. This analysis failed to detect any significant global difference in expression of these genes between males and females, regardless of the legs (Supplementary figure 5).

**Figure 5:**
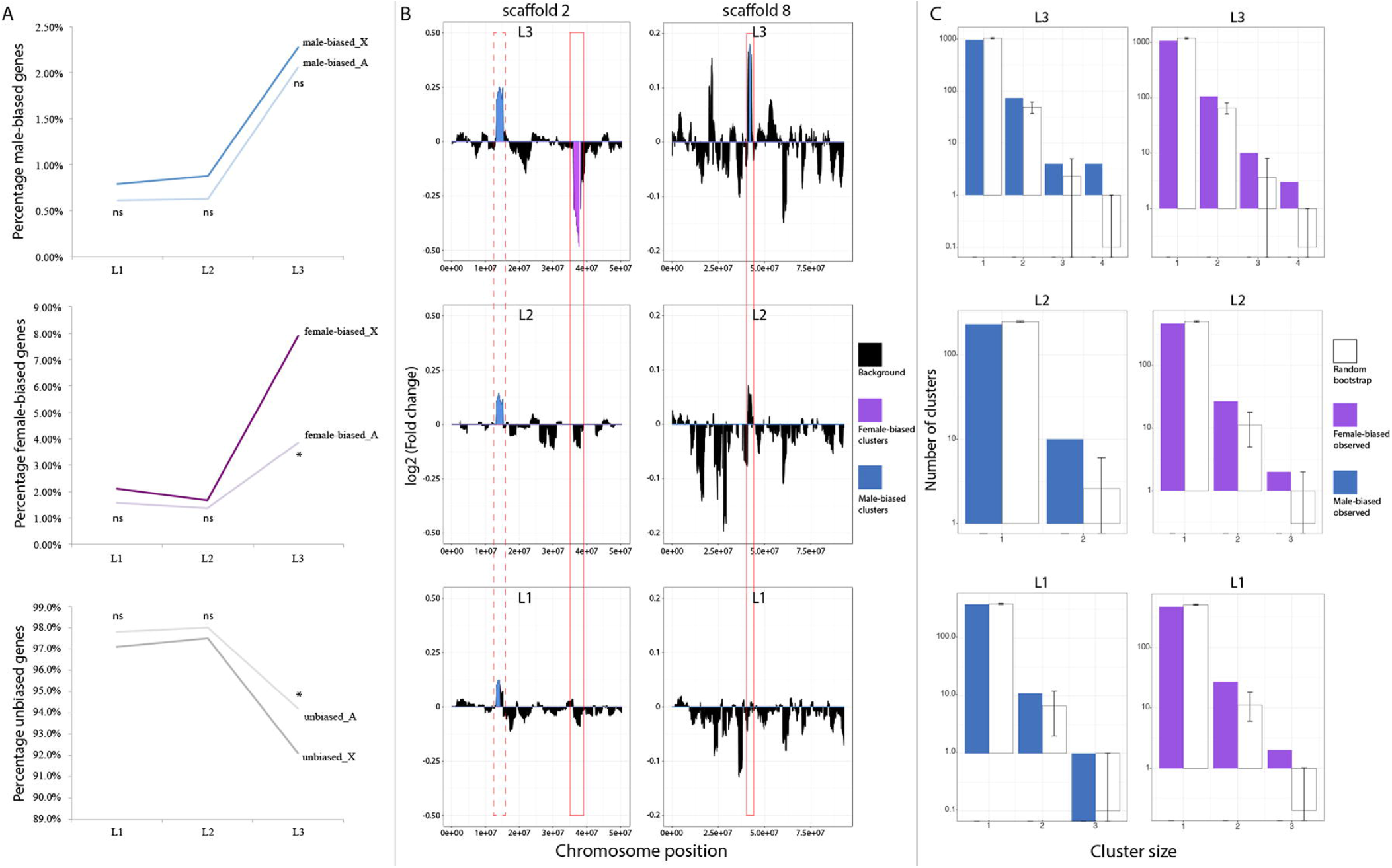
Genomic distribution of sex-biased genes in *M. longipes*. (**A**) Percentage of male-biased, female-biased and unbiased genes (from top to bottom) in the X chromosome and the autosomes across the three pairs of legs. Biased-distribution of the sex-biased genes on the X chromosome was estimated using the Fisher’s exact tests: * p-value< 0.05. (**B**) Large genomic clusters of sex-biased genes along scaffold #2 and the scaffold #8. Clusters highlighted in blue represent genomic regions enriched in male-biased genes. Clusters highlighted in purple represent genomic regions enriched in female-biased genes. Solid red frame indicates genomic clusters enriched in male- and female-biased genes specifically in the third legs. Dotted red frame indicates genomic clusters enriched in male-biased genes in all three legs but with different degrees of fold-change, recapitulating the degree of leg length exaggeration. (**C**) Genomic clusters of consecutive male- (blue) or female-biased (purple) genes in the three pairs of legs. Cluster size indicates the number of consecutive genes. Note that the y axis is log scaled. Error bars indicate fluctuation intervals.

Another potential effect of sexual selection on genome architecture is through rearrangements of genes or large genomic regions within chromosomes ^35–40^. We therefore performed a fine-scale visualization of the sex-biased genes along the thirteen largest scaffolds covering *M. longipes* genome (Figure 5B; Supplementary figure 6). We recovered few large genomic regions of 2 Mb significantly enriched in sex-biased genes (Supplementary figure 6; but see Material and methods). These include a total of 100 sex-biased genes (about 2% of the total number of sex-biased genes in all three legs), indicating that only a fraction of sex-biased genes arranges in such large genomic regions (Figure 5B; Supplementary figure 6). Among these, three large enriched regions located on scaffolds #2 and #8 contained a total of 36 sex-biased genes in the third legs (11 female-biased and 25 male-biased) (Figure 5B). Interestingly, two of these regions were specific to the third leg whereas the third indicated an enrichment of male-biased genes that was common to the three pairs of legs but with a higher degree of differential expression in the third legs (Figure 5B). In these regions, we could notably identify several unknown genes (10 out of 36 genes) including a cluster of four that were all strongly male-biased. Protein motif prediction, using Pfam, revealed a conserved domain of several transmembrane motifs in these four protein-coding genes.

Finally, we looked for small clusters of consecutive genes with similar patterns of expression in an attempt to assess common regulation ^41^. We found that over 15% of male-biased and over 20% of female-biased genes arrange in clusters of two to four genes in the third legs, while only about 8.5% and 10% respectively are expected under a null hypothesis of random gene order (p-value < 0.05; see material and method) (Figure 5C; Supplementary figure 7). More specifically, we found up to seven clusters of four consecutive sex-biased genes in the third legs while only a maximum of two of them were expected by random permutation (Figure 5C). In the second pair of legs, we also found that about 10% of the male- and female-biased genes are arranged in clusters of at least two genes, while 2 to 3% were expected by random permutation (p-value < 0.05; Figure 5C; Supplementary figure 7). In comparison with the third legs, clusters of male- and female-biased genes did not exceed two and three consecutive genes, respectively (Figure 5C). Male-biased genes in the first legs did not show any enrichment in clusters, and only one such cluster of three genes was detected (p-value > 0.05; Figure 5C; Supplementary figure 7). However, we found an enrichment of female-biased gene clusters, including 2 clusters of 3 consecutive genes (Figure 5C; p-value < 0.05; Supplementary figure 7).

### Molecular function of sex-biased genes

Finally, we aimed to determine the molecular function of the sex-biased genes in our dataset. Gene ontology (GO) term analyses revealed enrichment in translation, metabolic processes and Wnt signaling pathways for the male-biased genes in the third legs (Supplementary table 3). The “translation” GO term uncovered enrichment for several ribosomal proteins also known to play an essential role in cell proliferation in response to ribosomal stress ^42^. We also identified enrichment in molecular functions such as transferase activity indicative of possible post-transcriptional regulation differences between the two sexes. Female-biased genes in the third legs were enriched in various functions such as transcription factor, kinase, or GTPase activities that are probably involved in regulating biological processes such as transcription, metabolism, or signal transduction (Supplementary table 3).

## Discussion

Uncovering the genomic regulation underlying the process of phenotypic divergence between males and females is central to our understanding of morphological evolution ^5,8,30,43^. As an emerging model, *Microvelia longipes* offers exciting life history and ease of experimental manipulation to study how sexual selection can drive morphological and genomic adaptation ^22^ (also see companion paper Toubiana et al.). In a previous study, we found that the evolution of male third leg exaggeration was associated with intense competition between conspecific males to dominate egg-laying sites ^22^. The current study sheds light on the regulatory processes, both developmental and genomic, underlying this sex-specific exaggeration.

We identified a signature of leg exaggeration among sex-biased genes. Consistent with studies of sex-biased gene expression ^5,44–46^, we found that the degree of sexual dimorphism is associated with different patterns of expression among these genes (Figure 2&3). In our dataset, the most exaggerated legs mobilized more differentially expressed genes between the sexes and a higher degree of differential expression, especially in male-biased genes. Consistent with their potential role in driving the evolution of sexual dimorphism, male-biased genes were also relatively fast-evolving in comparison to female-biased and unbiased genes. This pattern is however common to all three legs, raising the question of pleiotropy in constraining adaptive evolution between the sexes. We found that a large proportion of sex-biased genes, especially in the third legs, displayed also tissue-specific expression (Figure 3A). Along with other studies showing less pleiotropy for sex-biased than unbiased genes ^47,48^, our results point out modularity as possible regulatory mechanism whereby tissues can evolve biased expression freely and acquire sex-specific phenotypes with little deleterious effects.

Comparing the three pairs of legs in *M. longipes* offers a unique opportunity to understand the genetic mechanisms underlying sex-specific structures as they present different types and degrees of sexual dimorphism, likely reflecting distinct selective pressures ^22^; (Toubiana et al. companion paper). For example, the development of the sex-combs, which are known to be under stabilizing selection ^49,50^, occurs in the first legs of males during the 5^th^ instar ^51^. In contrast, the exaggerated growth of male third legs, which also occurs during the 5^th^ nymphal instar (see companion paper Toubiana et al.), is an example of directional sexual selection that is absent, or at least strongly reduced, in the two other male legs ^22^. We therefore used *M. longipes* legs as comparison to link the development of various sexual dimorphisms with the regulation and genomic location of sex-biased genes in the three pairs of legs. The X chromosome, for example, has been hypothesized to be a genomic hotspot for sexual selection where female beneficial dominant mutations and male beneficial recessive mutations are expected to accumulate ^30,32,33^. However, interpreting the representation of sex-biased genes on the X chromosome is often influenced by dosage compensation ^4,32,33^. In *Drosophila* for example, the scarcity of male-biased genes on the X chromosome was suggested to result, at least partially, from dosage compensation. Our analyses detected a significant enrichment of the X chromosome with female-biased genes in the third legs, but not male-biased genes. This suggests that dosage compensation also operates in all legs of *M. longipes*, and is therefore unlikely to be responsible for the enrichment observed. Therefore, it is possible that this ‘feminized’ X chromosome represents a mechanism for the resolution of sexual conflict during the evolution of extreme sexual dimorphism in *M. longipes* third legs.

Previous studies have reported large genomic regions and profound genomic rearrangements (e.g. large chromosome inversions) in association with sexually dimorphic characters ^35,37–40^. In contrast, we found relatively few large but many small clusters of sex-biased genes in *M. longipes* genome (Figure 5). Moreover, these small and large enriched regions seem to be associated with the extreme elongation of male third legs, in terms of number, specificity or degree of differential expression, and may highlight some important genes and regulatory processes involved in sex-specific trait exaggeration. It is also important to note that previous studies on genomic clusters of sex-biased genes were primarily conducted on primary sexual organs, such as ovaries and testes ^35–40^. These tissues are highly complex, often express more sex-biased genes than secondary sexual traits and their evolution is considered to be under natural selection ^1,45,52–54^. Moreover, analyzing gene expression in these adult tissues does not capture the sex differences that are established during development. In the case of ontogenetic sexual dimorphism, it is expected that sexual selection will act on developmental regulatory processes ^5,11,55,56^. In this regard, our results offer the opportunity to test more accurately the role of sexual selection on gene and genome evolution by directly linking the development of sexual dimorphism with patterns and genomic locations of sex-biased genes in the three pairs of legs.

## Material and methods

### Population sampling and culture

A *Microvelia longipes* population was collected during fieldwork in French Guyana in Crique Patate near Cayenne. The bugs were maintained in the laboratory at 25°C and 50% humidity in water tanks and fed on crickets. Inbred populations were generated as described in ^22^.

### Statistics and leg measurements

All statistical analyses were performed in RStudio 0.99.486. For the PCA analysis on leg length, we used twenty males and females from each inbred population and measured them with a SteREO Discovery V12 (Zeiss) using the Zen software.

### Sample collection, assembly and annotation of the *M. longipes* genome

Hundreds of individuals (males and females mixed) were collected from three inbred populations and frozen in liquid nitrogen before DNA extraction. Genomic DNA was extracted and purified using the Genomic-tip 20/G DNA extraction kit from Qiagen. Genome sequencing, using a mix of Illumina mate pairs and PacBio libraries, was performed at the Beijing Genomics Institute. Chromosome-length scaffold assembly was performed by Dovetail Genomics using Hi-C/Hi-Rise libraries. Supplementary table 1 summarizes the sequencing strategy employed.

The genome sequence was polished using Illumina libraries (Supplementary table 1) and Pilon ^57^. Three different automatic annotation strategies, namely Braker, Maker and StringTie were tested to annotate the genome ^58–60^. These annotations were based on the leg transcriptomic dataset generated in this study (36 samples in total), a transcriptome from whole-body individuals collected at all developmental stages (1 sample) and a transcriptome from a third inbred population not mentioned in this study (18 samples). Braker and Maker pipelines also performed de novo automatic annotations. Maker and Stringtie annotations yielded lower BUSCO quality and manual quality assessment using JBrowse revealed a relatively high number of gene fragmentations that were poorly supported by the alignments. We therefore used Braker annotation for further analyses (Supplementary figure 1).

For Braker annotation, we used Hisat2 alignment files from each transcriptomic sample to train Augustus with UTR option. Final annotation includes 26,130 genes and 27,553 transcripts.

### Sample collection and preparation RNA-sequencing

We collected leg tissues from male and female 5th nymphal instars (two days after molting within a six hour time window) that belonged to two inbred populations that differ in average size (see ^22^). All individuals were raised in the same laboratory condition and fed with nine fresh crickets every day until the 5th instar. Individuals from the same inbred population were raised in the same water tank. The three replicates of each condition (lines, sexes and legs) correspond to a pool of 20 individuals chosen randomly (Supplementary figure 2). The dissection of the three pairs of legs, dissociated from the thorax, was performed in RNAlater (Sigma) using fine needles; each pair of legs was incubated immediately on ice in tubes filled with TRIzol (Invitrogen). RNA extractions were performed according to manufacturer protocol. The concentrations were assessed using the Qubit 2.0 Fluorometer (Invitrogen). Quality of RNA samples, library construction and sequencing were performed by Beijing Genomics Institute. The samples were sequenced using HiseqXten sequencing technology with a paired-end read length of 150bp.

### Transcriptome assembly, mapping and normalization

Read quality was assessed with FASTQC version 0.10.1 (http://www.bioinformatics.babraham.ac.uk/projects/download.html), and trimmed with TRIMMO-MATIC version 0.32. Specifically, reads were trimmed if the sliding window average Phred score over four bases was <15 and only reads with a minimum length of 36bp were kept. Braker annotation was used as reference for read alignment and the transcriptome quantification. We obtained around 90% alignment rate on the genome and about 72% of uniquely mapped reads using Hisat2 method (Supplementary table 4). The latter condition was used for the estimation of transcript abundances and the creation of count tables (raw counts, FPKM and TPM tables) were performed using the StringTie pipeline (Supplementary table 5) ^60,61^. The abundance of reads per gene was finally calculated by adding the read counts of each predicted transcript isoforms.

### Comparative transcriptomics: analyses of variance

Initially the transcriptomic approach was performed on three levels of comparisons; namely the lines, the sexes and the legs (Supplementary figure 2). The first three axes of variation in gene expression explained 57.1% of the total variation and separated the two inbred populations (Figure 1D). This confirms the genetic similarity that exists between individuals of the same inbred population. In order to correctly assess the influence of sex and leg comparisons on gene expression variance, we corrected for the line effect using a Within-Class Analysis ^62^. After correction, the first major axis of variation separated male and female conditions, while PC3 explained the variation between legs (Figure 1E).

### Identification of sex-biased genes

We first filtered transcripts for which expression was lower than 2 FPKM in more than half of the samples after combining the two inbred populations (12 samples total). Transcripts with average expression that was lower than 2 FPKM in both males and females were also discarded. The number of reads per “gene” was used to identify differences in expression among the different conditions of interest using DESeq2 ^63^. Differential expression analyses between males and females were performed on the two lines combined as we aimed to identify genes involved in male third leg exaggeration, which is a common feature to both lines. We repeated the differential expression analyses in lines separately and generally found high expression similarities between the two lines (data not shown). The differential expression analysis was also corrected for the line effect and we called sex-biased genes any gene with a fold-change > 1.5 and a Padj < 0.05. Other differentially expressed genes that do not fit such criteria (e.g. fold-change > 0 and a Padj < 0.05) are specified in the appropriate section.

### Interaction between leg and sex regulations

In order to detect a possible interaction between leg and sex regulations, we combined our list of sex-biased genes with another list of genes that were identified as differentially expressed between legs of the same sex (i.e., leg-biased genes) (see companion paper Toubiana et al.). Using Fisher’s exact tests, we then identified possible enrichment of genes with both sex- and leg-biased expression among the genes expressed within each tissue.

We also used the interaction model implemented in DESeq2. For this, we first filtered for lowly expressed transcripts by removing all transcripts for which the expression was lower than 2 FPKM in more than two-thirds of the samples (36 samples total). This filtering process leaves 9364 transcripts for the differential expression analysis. The interaction model between legs and sexes, after correcting for the line effect, revealed 2 genes for the third and first leg comparison and no gene for the third and second leg or the second and first leg comparisons. When we looked at these two genes in the differential expression analyses without the interaction effect, we found that one of them (g7203) was detected as male-biased in the third leg but not leg-biased whereas the second gene (g23967) was both male-biased and upregulated in the third leg compared to the first.

### Hierarchical clustering

Average expressions of sex-biased genes in the different tissues were clustered using Euclidean clustering in the R package PVCLUST version 1.3-2 ^64^ with 1000 bootstrap resampling. Heatmaps and clustering were performed using the log2(TPM) average expression of each gene from each tissue. Heatmaps were generated using the R package GPLOTS version 3.0.1.1.

### Sex-biased gene distribution between chromosomes

#### Sex chromosome identification

Sex in the Gerromorpha is genetically determined and established by either the XX/XY or XX/X0 ^65,66^. In *M. longipes*, Illumina genomic sequencing containing only males was used to align genomic reads against *M. longipes* genome and extract the genomic coverage of each scaffold. The scaffold 1893 was the only scaffold among the 13 biggest scaffolds (more than 90% of the genome) that presented twice less coverage than the other scaffolds. To finally assess the identity of the X chromosome in *M. longipes*, we monitored the gene expression and found that the scaffold 1893 included both male- and female-biased genes, excluding this scaffold to be the Y chromosome. We also looked for a possible Y chromosome by identifying scaffolds with similar genomic coverage as the X chromosome but containing genes with only male-biased expression. We did not find any among the fifty largest scaffolds, suggesting that *M. longipes* has a XX/X0 sex determination system or presents a highly degraded Y chromosome.

#### Genomic distribution of sex-biased genes

We identified the genomic location of each gene and selected genes with a fold change superior to 1.5 between males and females as sex-biased genes (Padj < 0.05). Over- or under-representation of sex-biased genes in the X chromosome (scaffold 1893) compared to the autosomes (12 other largest scaffolds) was tested using Fisher’s exact tests.

#### Estimation of dosage compensation

To compare the average level of gene expression between males and females in the X chromosome we first selected expressed genes with FPKM > 2 in at least half of the samples (12 samples per leg). We also averaged gene expressions between replicates and lines before testing for differences in expression (Wilcoxon tests on the log2(FPKM)).

### Detection of large sex-biased gene regions

To detect large chromosomal regions enriched in sex-biased genes we developed a bootstrapping method based on sliding windows of 2 Mb with a step size of 100 kb (Supplementary figure 8). Gene density calculation revealed that on average, genes are found every 20 kb in *M. longipes* genome. This pattern was homogeneous among chromosomes (Supplementary table 6). We therefore split each chromosome into bins of 100 kb and generated sliding windows of 2 Mb (20 bins) to include approximately 100 genes per window in the analysis (Supplementary figure 8B, Supplementary table 6). We used two scaffolds, one scaffold with two enriched regions (scaffold 2) and a scaffold with no enriched region (scaffold 1914), to repeat the analysis with smaller regions (1 Mb, 500 kb, 250 kb and 120 kb). We found similar results in both scaffolds, regardless of the size of the region, indicating that our analysis is statistically robust and is not missing information.

#### Fold-change reassignment and gene position

From the DESeq2 analyses, all expressed genes were associated with a log2 fold change (Log2FC) and a p-value (Padj). Unexpressed genes (FPKM < 2) were assigned a log2FC of 0 and a p-value of 1. Among the expressed genes, we switched the log2FC to 0 for the unbiased genes (Padj > 0.05), in order to directly assess sex-biased genes based on log2FC values (Supplementary figure 8A).

In a second step, we merged the dataset on sex-biased expression with the gene positions (Supplementary figure 8A).

#### Genome-wide detection of sex-biased gene regions

A mean log2FC was calculated for each window and reported along the chromosomes to reveal genome-wide regions of sex-biased genes (Supplementary figure 8B).

#### Bootstrapping method

To test whether these regions are significantly enriched in male or female-biased genes, we developed a bootstrap approach (Supplementary figure 8C). As the mean expression level of a gene influences the log2FC value (i.e. genes with low expressions are more likely to have high log2FC values and genes with high expression are more likely to be differentially expressed), we created 5 categories of genes based on their expression levels (baseMean values from DESeq2 tables). We then reassigned randomly, within each category, the log2FC at each gene position in the genome. This step was performed 100 000 times, therefore generating 100 000 random log2FC profiles.

Finally, to test for the significant enrichment of gene in these regions, we compared for each bin the observed log2FC values with the log2FC values generated from the bootstrap. To call for significantly enriched region of sex-biased genes, we identified regions for which the observed log2FC value was higher (male-biased) or lower (female-biased) than the 97500 (out of 100 000) bootstrap values generated randomly (Supplementary figure 8D). We finally applied a Bonferroni correction, correcting the bootstrap values by the total number of independent windows in the genome (n=300), leading to a bootstrap threshold of 99992. We note that this analysis is rather stringent (possibly missing other important enriched regions), but the highlighted regions are unambiguous (low sensitivity but high robustness).

### Detection of clusters of consecutive sex-biased genes

This analysis was primarily inspired from Boutanaev et al. ^35^. In short, we determined clusters by ordering genes along the genome and detecting regions of consecutive male- or female-biased genes (Padj < 0.05). To avoid identifying clusters overlapping two different chromosomes, we performed this analysis on the thirteen largest scaffolds separately. We then tested whether the observed distribution of genes differed from a stochastic distribution by randomly assigning a genomic position to unbiased, male-biased and female-biased genes respectively. The proportion of sex-biased genes found in clusters as well as the distribution of cluster sizes was calculated by averaging 1000 iterations (Supplementary figure 8). P-values were extracted from the 95% fluctuation intervals calculated from the 1000 randomized iterations.

### Gene Ontology analysis

Gene names and functions were annotated by sequence similarity against the NCBI ‘non redundant’ protein database using Blast2GO. The Blast2GO annotation was then provided to detect Gene Ontology terms enrichment (p-value < 0.05) using the default method of TopGO R package version 2.34.0.

## Acknowledgements

We thank Jean-Nicolas Volff comments on the manuscript, Laurent Duret for help with dn/ds analyses and François Leulier, Kevin Parsons and Gaël Yvert for helpful discussions. This work was funded by ERC-CoG WaterWalking #616346 and Labex CEBA to A.K., a Ph.D. fellowship from *Ecole Doctorale* BMIC de Lyon to W.T, and Swiss National Foundation fellowship to R.A.

