## Supplementary material for "Impact of trait exaggeration on sex-biased gene expression and genome architecture in a water strider": Suplemental figures

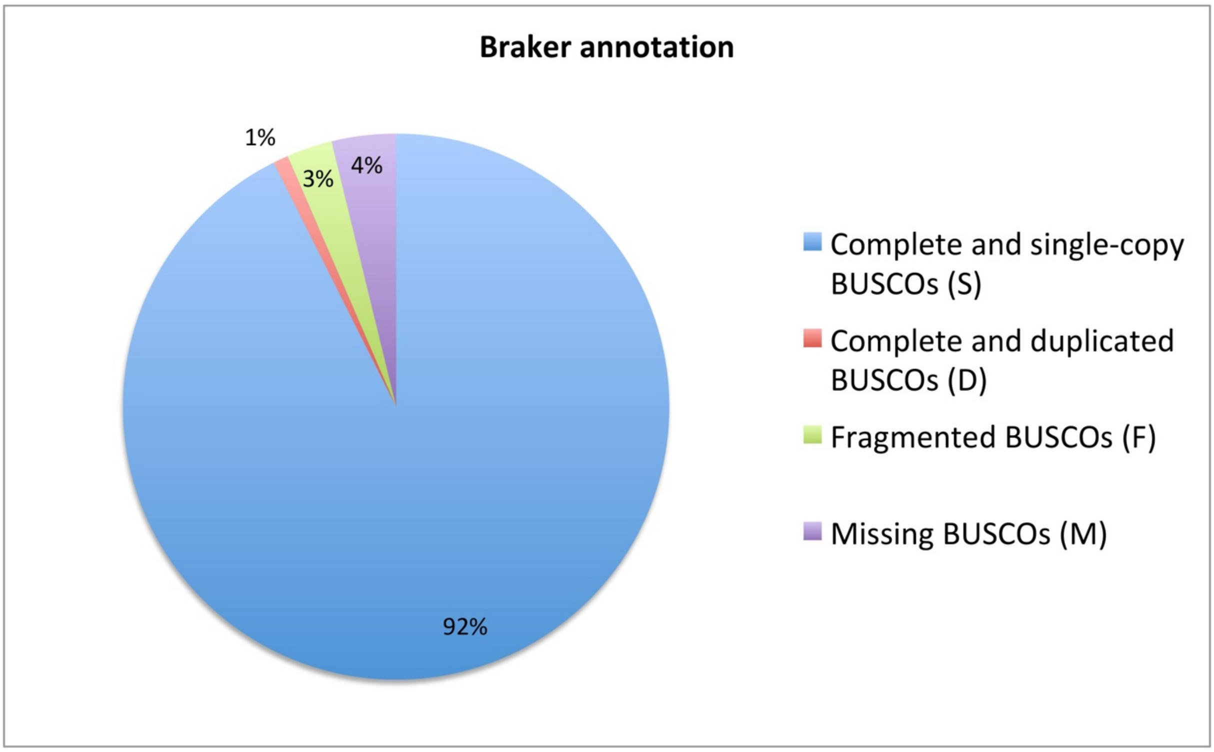


**Supplementary figure 1:** Diagram of BUSCO analysis run for the 26130 genes identified from Braker annotation.

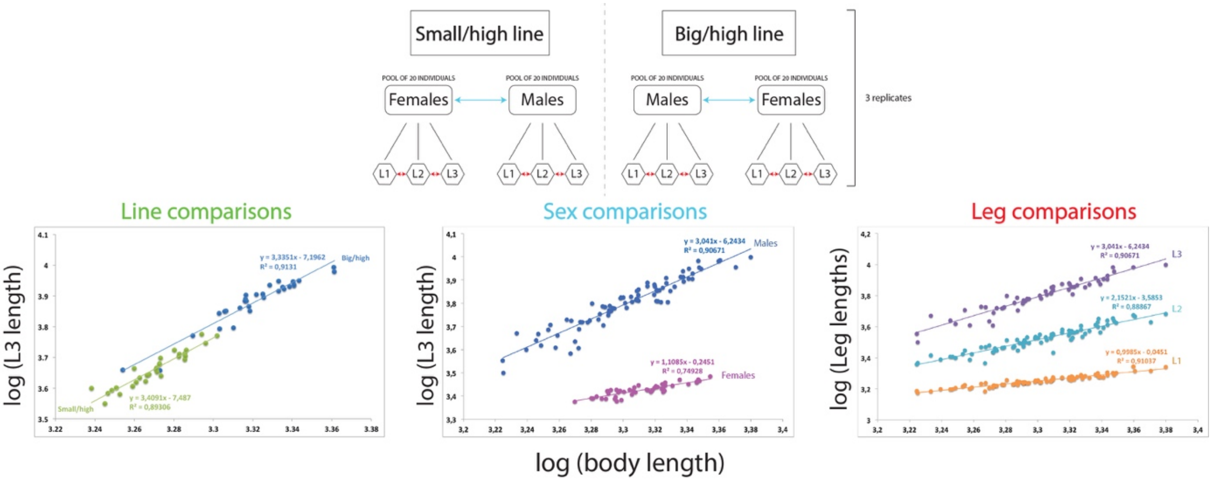


**Supplementary figure 2:** Experimental design of the comparative transcriptomic analysis.

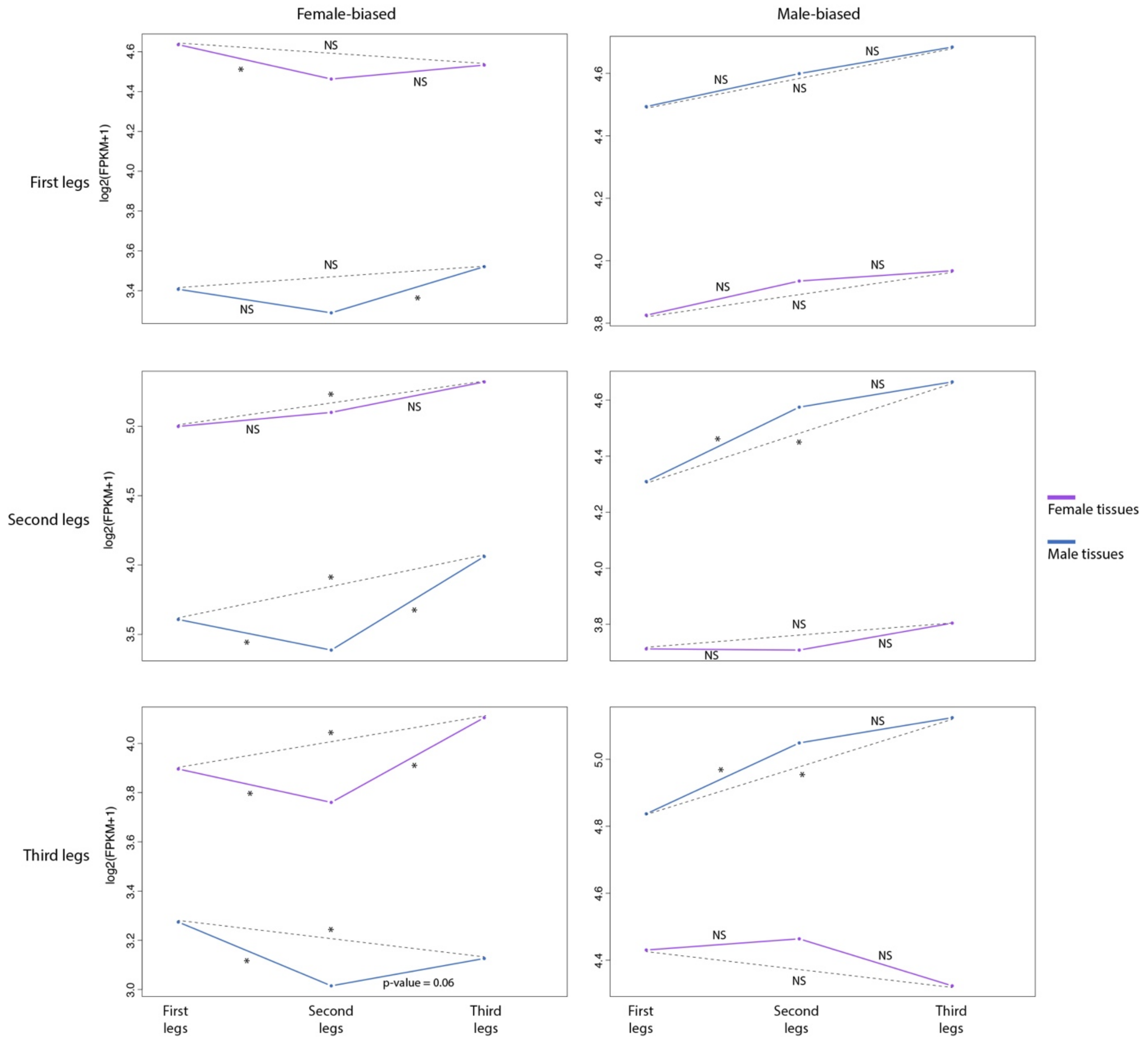


**Supplementary figure 3:** Expression (log2FPKM+1) of sex-biased genes in males and females across legs.

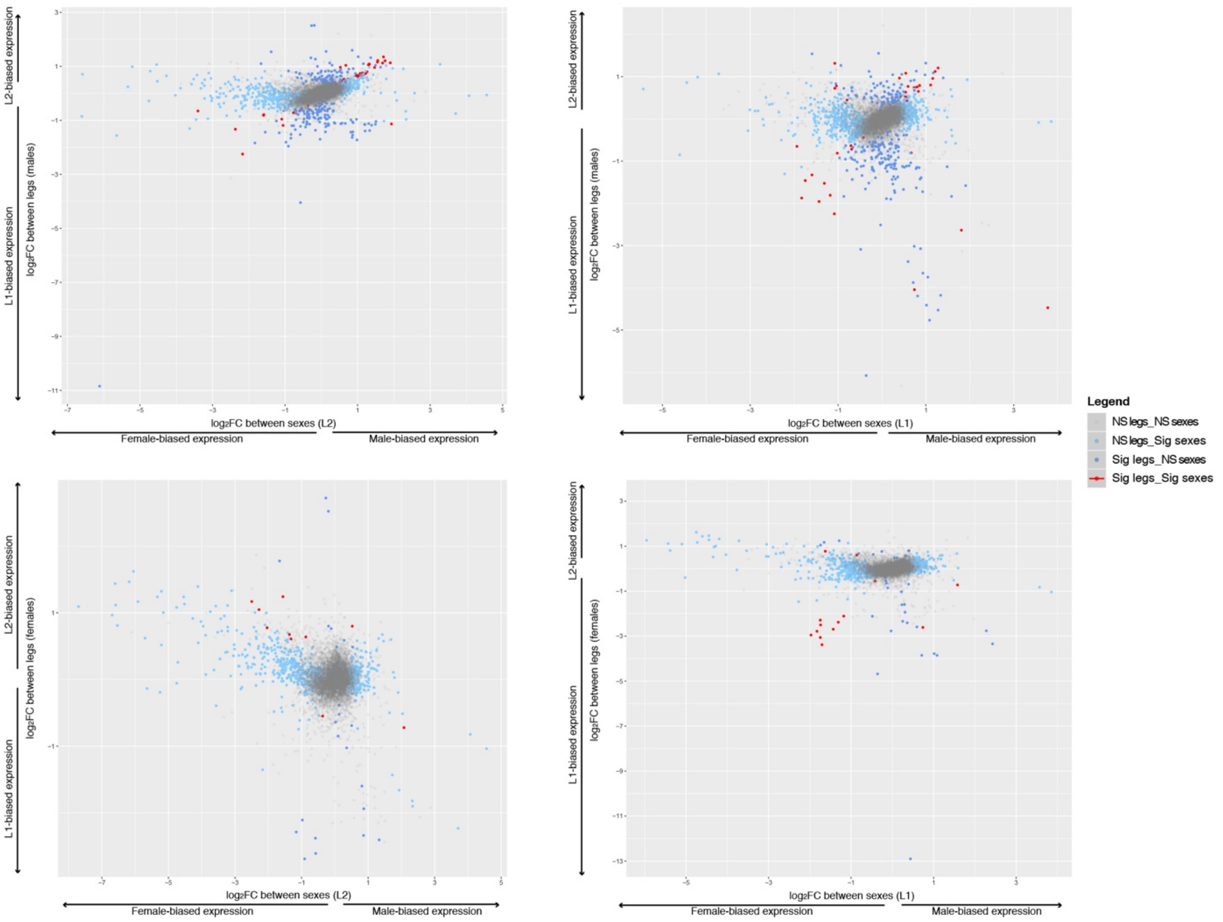


**Supplementary figure 4:** Correlation between leg- and sex-biased genes using the second legs as reference.

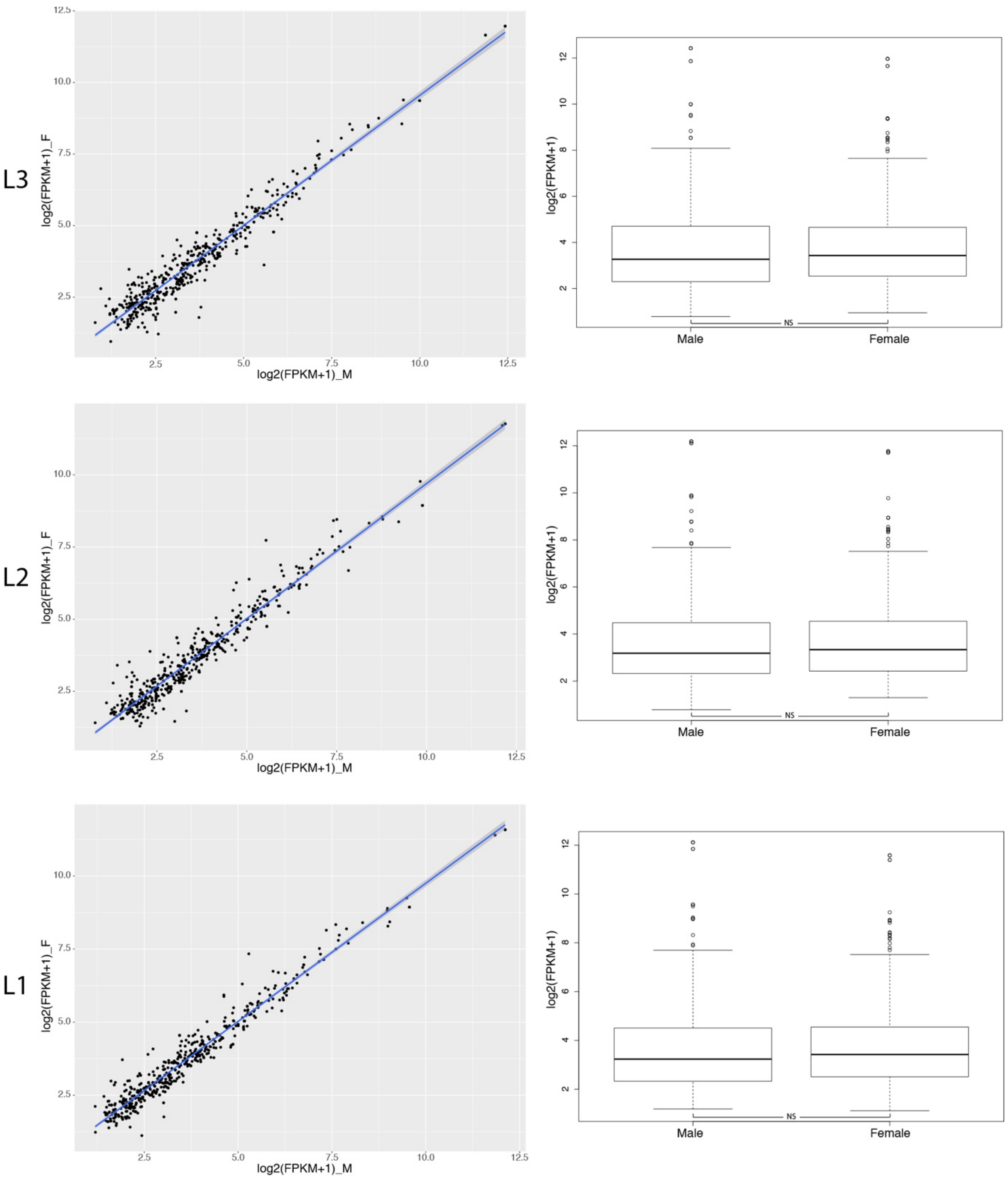


**Supplementary figure 5:** Average gene expression differences between males and females on the X chromosome. Regressions were fitted from a linear model.

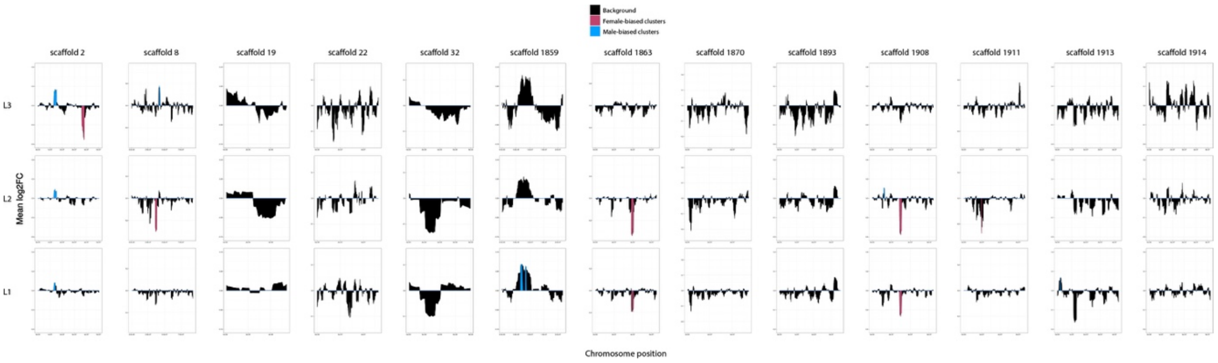


**Supplementary figure 6:** Genome-wide characterization of large genomic clusters

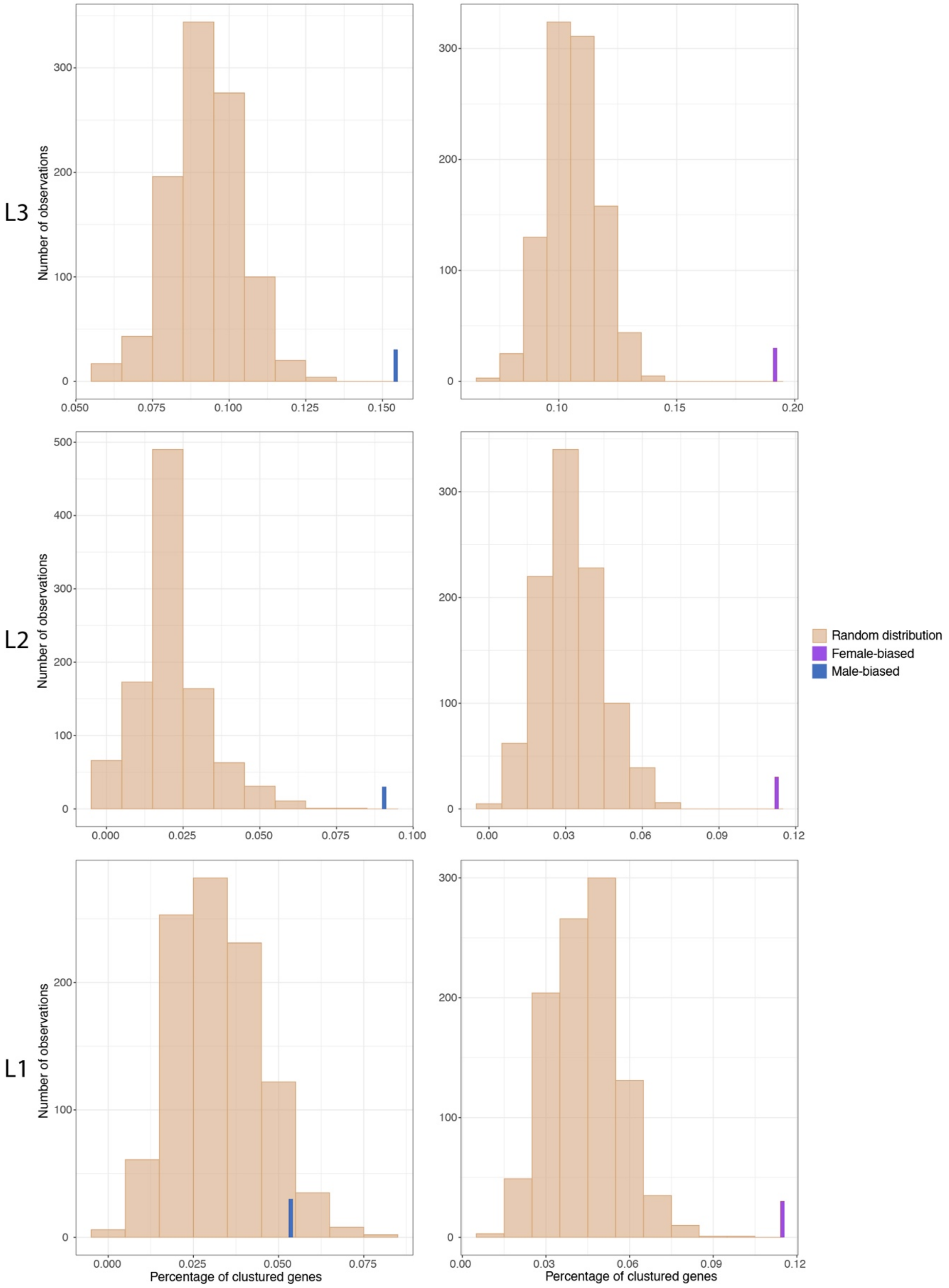


**Supplementary figure 7:** Distributions of the proportion of clustered sex-biased genes (at least two consecutive sex-biased genes) generated through 1000 random iterations in each leg separately. Blue and purple bars correspond to the observed proportions of male- and female-biased genes, respectively, in the three legs.

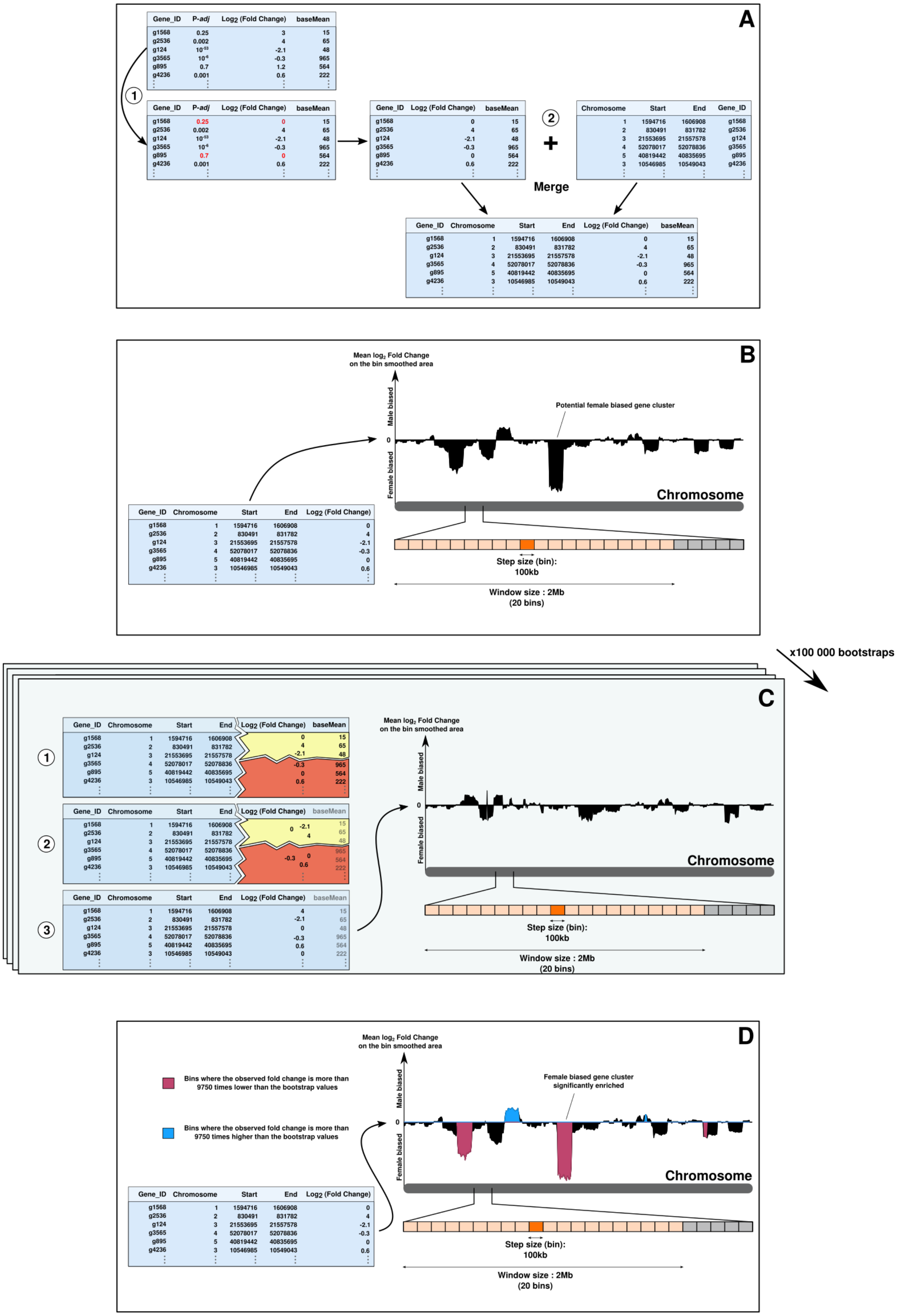


**Supplementary figure 8:** General pipeline of the bootstrap analysis to detect large genomic clusters of sex-biased genes.

**Supplementary table 1:** Genomic libraries used to sequence and assemble *M. longipes* genome. Big/low, Big/high and Small/high being the three inbred populations used for the genome sequencing.

**Supplementary table 2:** GLM statistics for dN/dS analyses.

**Supplementary table 3:** Gene Ontology (GO) terms for male and female-biased genes in the three legs.

**Supplementary table 4:** Transcriptome metrics with the number of reads per library and the alignment rate of these reads on the *M. longipes* genome.

**Supplementary table 5:** Summarizing tables on raw read counts, FPKM counts, blast analysis and the lists of sex-biased genes across legs.

**Supplementary table 6:** Summary table of the average gene density per scaffold.
